# Asymmetric co-substrate usage at a metabolic branch point as a minimal network motif that can drive overflow metabolism

**DOI:** 10.1101/2025.06.05.658108

**Authors:** Robert West, Sonal, Tailise D.S.G. Rodrigues, Wenying Shou, Orkun S. Soyer

## Abstract

Metabolic overflow is pervasive from microbes to tumors: as glucose consumption increases, cells switch from respiration to respirofermentation with excretion of organic acids or ethanol. How do cells regulate their fluxes to implement this switch? Using simplified models of a branch point, we found that three independent “built-in” mechanisms can each enable flux switching at a branch point: differential enzyme kinetics at the two branches, allosteric regulation of a branch enzyme, and asymmetric co-substrate (e.g. NADH/NAD^+^) usage across the two branch point reactions. Notably, the co-substrate mechanism achieves several fold higher sensitivity, which can be further enhanced by the other two mechanisms. We then applied this theory to the yeast pyruvate branch point underpinning overflow metabolism by developing a compartmentalized model with or without NADH dynamics in cytoplasm and in mitochondria. Crucially, only models that included NADH dynamics recapitulated experimental data: (a) NADH level increases nonlinearly with increasing glucose influx; (b) this increase occurs at a lower glucose threshold than that for overflow; and (c) glucose threshold for overflow increases when NADH oxidase is expressed in mitochondria but not in cytosol. Thus, cosubstrates are not merely passive carriers of chemical groups; their asymmetric utilization at a branch point offers an intrinsic, network-embedded regulatory layer that synergises with enzyme-based control. Beyond explaining overflow metabolism in yeast, this theory offers a metabolic engineering strategy for re-routing flux through manipulating co-substrates.

## INTRODUCTION

Overflow metabolism, also known as aerobic glycolysis, ‘Warburg’ or ‘Crabtree’ effect, is a broadly observed cellular phenomenon, seen in tumor cells, yeast, and bacteria^1–7^. It involves an increased flux into fermentation products, such as lactate, ethanol, and acetate, with increasing glucose consumption rates, despite aerobic conditions and continued respiration (respiro-fermentation)^2,6^. Since respiration produces more ATP per glucose compared to fermentation, and since cells achieve a higher growth rate under respiro-fermentation, metabolic overflow is seen as cells trading-off efficiency in energy harvesting for faster growth^6,9^.

Despite its importance and seemingly universal presence, the causes of overflow metabolism are still unclear. At a phenomenolgical level, overflow metabolism is explained as arising from cellular constraints^10^. Yet, the nature of such a constraint is debated. For instance, several modelling studies proposed that protein synthesis can become a constraint under fast growth conditions, and that cells switch to fermentation due to the lower protein cost of fermentation compared to respiration^1,11^. However, proteomics measurements are inconclusive on the extent of changes in pathway-specific enzyme levels^12–14^. In addition, recent experimental studies report contrasting results on protein efficiency of fermentation vs. respiration^15,16^. Importantly, metabolic overflow can happen within a few minutes^4,17^, and similarly, other metabolic dynamics can be fast and cannot entirely be explained by changing enzyme expression levels^14,17–20^. Thus, other mechanisms based on, for example, allosteric regulation or metabolic network structure, are likely to be involved.

Within central metabolism, metabolic overflow specifically relates to the pyruvate branch point (Fig. 1A): Pyruvate is converted either toward organic acids via fermentation or toward acetyl-CoA, which feeds the tricarboxylic acid (TCA) cycle and respiration. In principle, branch points are well-suited for regulation. As shown in a seminal modelling study, the flux distribution between two branches can be altered by modulating the levels and activities of the branch-specific enzymes^21^. Indeed, for the pyruvate branch point, enzymes immediately downstream of the branch point, including pyruvate decarboxylase (PDC) on the fermentation branch and pyruvate dehydrogenase (PDH) on the respiration branch, are subject to extensive regulation^22–24^. In mammalian cells, for example, overflow metabolism can be influenced by altering the phosphorylation state of PDH via its kinase^25^. Additionally, the branch point enzyme PDC is known to be allosterically activated by its substrate pyruvate^24^. There are also allosteric regulations of upstream enzymes in glycolysis, such as the pyruvate kinase (PYK) and phosphofructokinase (PFK), that are implicated in the regulation of the glycolysis-gluconeogenesis switch^26,27^.

**Figure 1:**
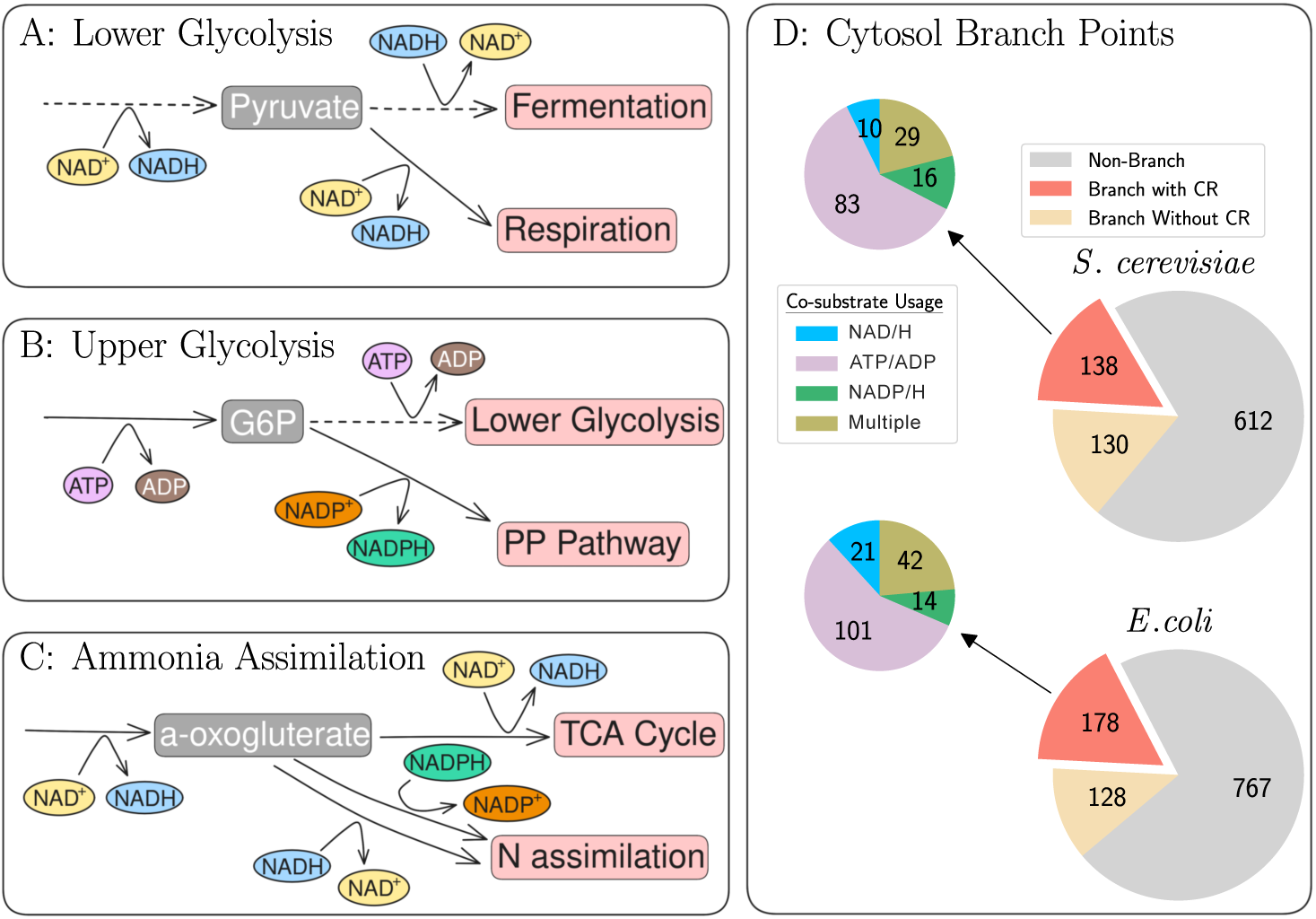
Branch points in central metabolism. (A-C) Cartoon representation of different examples of branch points with co-substrate-involving reactions (CRs). Each cartoon is a simplification of the corresponding reaction maps included in the Yeast Pathways database^54^. In particular, dashed arrows indicate multiple reactions, grey and pink boxes indicate metabolites and processes respectively, and other colours indicate different co-substrates. EC Numbers and pathways used to create the cartoons are: (A) Lower Glycolysis - 1.2.12 (glycolysis I), 1.1.1.1 (superpathway of glucose fermentation), 1.1.1.41/1.1.1.286 (TCA cycle, aerobic respiration). (B) Upper Glycolysis - 2.7.1.1/2.7.1.2 (superpathway of glucose fermentation), 2.7.1.11 (glycolysis I, from glucose 6-phosphate), 1.1.1.49 (pentose phosphate pathway, oxidative branch I). (C) Ammonia Assimilation - 1.1.1.41/1.1.1.286, 1.2.1.- (both from TCA cycle and aerobic respiration), 1.4.1.4 (glutamate biosynthesis from ammonia), 1.4.1.14 (L-glutamate biosynthesis IV). (D) Branch-point metabolites using specified co-substrates in *S. cerevisiae* and *E. coli*. Genome-scale models for *S. cerevisiae* ^36^ and for *E. coli* ^37^ are analysed for metabolites that are consumed in two or more reactions, excluding excretion reactions. All these metabolites are counted as ”branch-point metabolites” and are further analysed for the presence of immediate consuming reactions involving specific co-substrates. Analysis is presented for metabolites in the cytosol with co-substrates ATP and NAD(P)H (and their different forms), as these accounted for most of the cases (see also *Methods*).

The pyruvate branch point, like many other branch points in central metabolism (Fig. 1), features reactions dependent on co-substrates — metabolites that act as donors and acceptors of chemical groups. Examples of co-substrates include electron carriers NAD^+^/NADH and phosphate carriers ATP/ADP. While metabolic models commonly assume constant co-substrate concentrations, relaxing this assumption reveals that co-substrate dynamics can become flux limiting^28–32^. Experimentally, modulating NADH dynamics (via introducing enzymes that consume or produce NADH) changed the overflow initiation point in yeast and bacteria^33,34^, and similarly altered metabolic overflow in mammalian cells^25,35^. These experimental findings lead to the hypothesis that NADH dynamics are directly interlinked with the rerouting of flux at the pyruvate branch during metabolic overflow.

Here, we ask whether network architecture itself, through co-substrate dynamics, can regulate fluxes. We address this question using minimalistic models of branch points with or without allosteric regulation and co-substrate usage. In these models, we systematically vary the placement of co-substrate reactions. We find that asymmetric co-substrate usage at the branch point enables, on its own, a form of ‘self-regulation’: As influx into the upstream pathway increases, the flux distribution across the two branches can undergo an abrupt switch. While we find a similar response with differential enzyme kinetics or allostery at the branch point, the co-substrate-based mechanism offers greater parameter flexibility and sensitivity than these other mechanisms, and can synergize with them. To apply this theory to a realistic case, we develop a model of the pyruvate branch point in yeast, accounting for NADH dynamics across the cytosol and mitochondria, as well as incorporating known enzyme kinetic parameters and allosteric regulation of PDC. We find that both allosteric regulation and NADH dynamics can separately capture the experimentally-observed nonlinear flux switching with increasing glucose influx. However, only when NADH dynamics are included can the model qualitatively capture additional experimental observations: 1) NADH fraction undergoes an upward switch with increasing glucose influx, 2) the NADH switch occurs at a *lower* glucose influx than the overflow switch^5,19^, and 3) perturbing NADH turnover alters the glucose influx threshold for initiating ethanol overflow^33,34^. We conclude that asymmetric co-substrate usage at branch points can introduce a self-regulatory flux switching mechanism that operates alongside mechanisms such as allosteric regulation to drive overflow metabolism. This theory suggests that modulating co-substrate dynamics at branch points offers a general principle for explaining and engineering diverse metabolic behaviors.

## RESULTS

### Co-substrate reactions (CRs) are commonly found at or around branch points in cell metabolism

Branch points and co-substrate reactions (CRs) are common throughout cellular metabolism (Fig. 1). Interestingly, different branch points show different placement of CRs at and around these junctions. For example, the pyruvate branch point in yeast harbours three NADH-linked CRs, one immediately downstream of the branch point, and another two some distance upstream or downstream (Fig. 1A). In other examples, upper glycolysis features a branchpoint involving CRs that utilize either ATP or NADP^+^ (Fig. 1B), while the ammonia assimilation branch point features CRs involving NADH and NADPH (Fig. 1C). Using the latest genome scale models, we have identified hundreds of metabolic branch points in *Saccharomyces cerevisiae* and *Escherichia coli* ^36,37^, over 50% of which utilize co-substrates (Fig. 1D).

### Differential enzyme kinetics on the two branches and allostery can generate weak flux-switching responses to increasing pathway flux

How might the placement of CRs around a branch point influence flux distribution between the branches? We explore this question by first focusing on didactic, minimalist models of a branch point, i.e. branch-point motifs. We consider enzymatic reactions to be irreversible, but background co-substrate conversion is considered reversible to account for the co-substrate’s dynamic role as both donor and acceptor of a chemical group (e.g. electrons in the case of NADH/NAD^+^) (see *Methods* and Fig. S1). To understand the ”regulation potential” in a branch-point motif, we focus on steady state flux fraction: *F_f_* = *F*_upper_/(*F*_lower_+*F*_upper_) as a key variable describing branch-point flux dynamics, and consider how *F_f_* is affected as the upstream pathway influx *v*_in_ increases.

We compare *F_f_* across four scenarios (Fig. 2): with neither allosteric regulation nor co-substrate dynamics (panel A), with allosteric regulation of the upper-branch enzyme activity only (panel B), with co-substrate dynamics only (panel C), and with both allosteric regulation and co-substrate dynamics (panel D). For each scenario, we consider three cases for enzyme kinetics:

**Figure 2:**
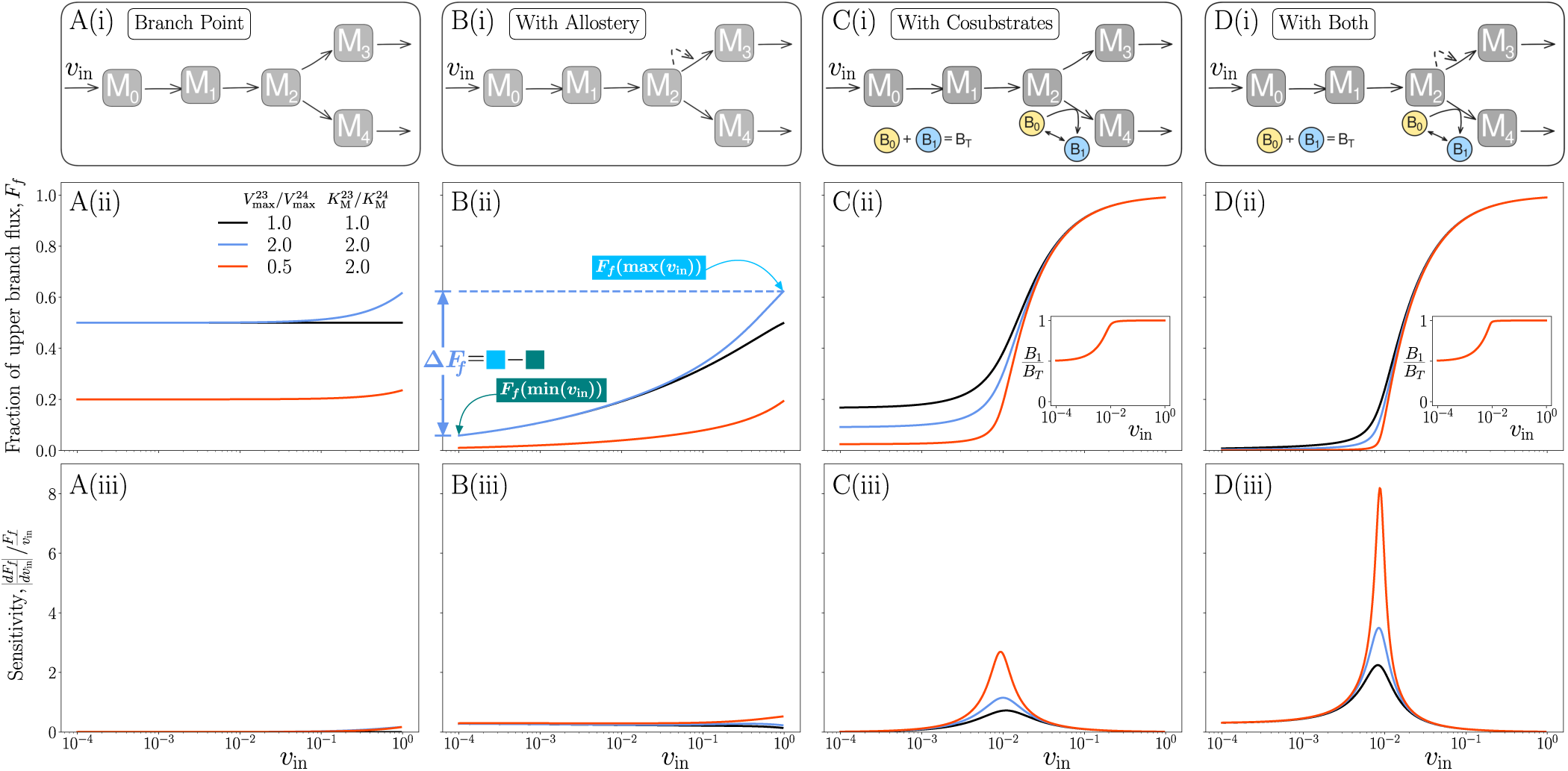
Co-substrate reactions synergise with allostery to enable flux switching with increasing influx. **(A-D)(i)** Simple branch-point motifs with different specifics: motif without co-substrates, but with (B) or without (A) allosteric activation of upper branch enzyme by its substrate; motif with a single co-substrate reaction on one branch (C) and with co-substrate and allostery (D). **(A-D)(ii):** Simulation results from models corresponding to the cases shown in (i). Each panel shows the steady-state flux fraction, *F_f_*, which is the ratio of steady state flux from *M*_2_ to *M*_3_ over total flux at the branch point, against increasing influx into the pathway *v*_in_ (see *Methods*). On panel B(ii), we visualize Δ*F_f_*, which gives the difference of flux fraction at the highest and the lowest *v*_in_, while insets in C & D show the *B*_1_/*B*_T_ against *v*_in_. **(A-D)(iii):** Sensitivity of the gradient of *F_f_* with respect to *v*_in_. All reactions, apart from the background conversion of *B*_0_ and *B*_1_, are modeled as irreversible. Different colored lines indicate the different enzyme kinetic parameters used, as shown on the legend of panel A and applies to all panels. All other parameters are set arbitrarily to 1. For panels C-D, parameters relating to *B* and its turnover are all set to 1, except for 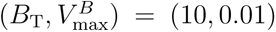. In panels B and D, we additionally have allosteric regulation of the *M*_2_ *M*_3_ reaction by its substrate *M*_2_. This is modeled by using (*m*_2_)*^n^* and 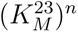 where *n* = 1.3 in the Michaelis-Menten equation for this reaction (see *Methods* and Fig.S1).

(i) upper and lower branch enzymes having the same kinetics (Fig. 2, black lines), (ii) the upper enzyme having weaker substrate affinity but higher maximum rate (Fig. 2, blue lines), and (iii) the upper enzyme being weaker in both maximum rate and substrate affinity (Fig. 2, red lines). Overall, our choices for these different scenarios are inspired by the pyruvate branch point, where PDC and PDH have differing kinetic parameters (Table S1), and where PDC is allosterically activated by its substrate, pyruvate.

For symmetric enzyme kinetics without co-substrate dynamics or allostery, unsurprisingly, the flux is split evenly between the two branches (*F_f_* = 0.5), and does not respond to an increase in influx *v*_in_ (Fig. 2A(ii), black line). When the two branch-point enzymes differ in maximal rate or substrate affinity, the flux distribution can shift away from an even-split and can show a weak response to the increase in influx *v*_in_(Fig. 2A(ii), blue and red lines). In this case, we have the following expression for *F_f_*, the flux fraction of the upper branch (metabolite 2 to 3 in Fig. 2A(ii)):

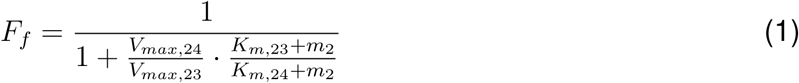

where *m*_2_ is the concentration of the branch point metabolite *M*_2_, and *V_max_* and *K_m_* represent the maximal rate and substrate affinity of the enzyme catalyzing the upper (subscript _23_) or lower (subscript _24_) branch. Thus, when the two enzymes have different substrate affinity, *F_f_* will respond to *m*_2_ which will in turn respond to influx *v*_in_ (Fig. 2A(ii), blue and red lines; see also Figure S2). Introducing allosteric regulation of the upper branch-point enzyme enhances this response further (Fig. 2B(ii)).

The magnitude of the *F_f_* response (Δ*F_f_*) can be quantified as the difference of flux fraction at the highest and the lowest influx levels (as illustrated in Fig. 2B(ii)), while its sensitivity (i.e. degree of response nonlinearity) can be measured by relative change in *F_f_* compared to that in *v*_in_. (Fig. 2A-D(iii)). This analysis shows that either differences in the kinetic parameters of branch-point enzymes or allosteric regulation of a branch-point enzyme can result in a response of *F_f_* to increasing influx, but these responses and their sensitivity are generally weak (Fig. 2A, B(ii)-(iii)).

### Co-substrate dynamics alone can facilitate flux switching in response to increasing pathway flux at a branch point

We now consider the case of placing a single co-substrate reaction (CR) immediately after the branch-point metabolite, utilizing one form of a co-substrate, B_0_ (representing for example NADH or NAD^+^) (Fig. 2C). For the simple motif shown in (Fig. 2C), we can again derive an expression for the flux distribution *F_f_* :

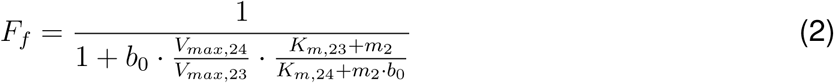

where the notation is as before and *b*_0_ represents the concentration of the *B*_0_ form of the co-substrate. As seen in Eq. 2, *F_f_* now depends on the concentrations of both *M*_2_ and *B*_0_, and this dependence does not cancel out even when the branch point enzymes have the same kinetics. Increasing *v*_in_ would in general lead to an increase in *m*_2_, which will increase B_0_ consumption and hence decrease *b*_0_ (Figure S2). Indeed, increasing *v*_in_ triggers flux switch to the upper branch (Fig. 2C(ii)), coinciding with a depletion of B_0_ (Fig. 2C(ii), inset).

Overall, as *v*_in_ increases, the lower branch becomes increasingly limited by the co-substrate B_0_ and hence the upper branch point gets increasingly more substrate (*M*_2_). This asymmetric co-substrate dependency between the two branch-point reactions drives the switch-like *F_f_* response to increasing *v*_in_.

The switch-like response in *F_f_* generated through co-substrate dynamics does not require specific kinetics or any allosteric regulation of the branch-point enzymes (compare same colored lines in Fig. 2A-C(ii)), and displays a higher sensitivity (compare Fig. 2A-C(iii)). The sensitivity of this response can be further enhanced by introducing differences in kinetic parameters and/or allosteric regulation (Fig. 2D(ii) and (iii)).

We conclude that introducing a single CR at the branch point creates an asymmetry arising from differences in co-substrate usage, which facilitates a switch-like response in *F_f_* at a threshold level of pathway influx, a form of self-regulation.

### Flux-switch dynamics are maintained over a wide range of co-substrate pool size and turnover rate

The switch-like behavior of *F_f_* at a threshold *v_in_*is driven by asymmetric co-substrate usage: while the CR branch experiences co-substrate depletion as influx increases, the non-CR branch is unaffected. Since co-substrate depletion can be influenced by the total co-substrate pool *B*_T_ and/or the maximum turnover rate 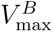, we examined how the magnitude of switch response Δ*F_f_* is influenced by co-substrate turnover dynamics.

Focusing first on the simplest branch-point motif with a CR (Figure 2C), we find that the large Δ*F_f_* response is maintained over multiple orders of magnitude of total pool size *B*_T_ and turnover rate 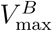 (Fig. 3A, see also Fig. S3). For this motif, large total pool size combined with slow turnover leads to a flux switch to the upper branch as influx increases (red region in Fig. 3A), while small *B*_T_ and high 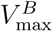 result in opposite switch direction, i.e. a negative Δ*F_f_* (blue region in Fig. 3A). This negative Δ*F_f_* dynamics can be understood by considering Eq. 2, and the differential effect of increasing *v*_in_ on *m*_2_ and *b*_0_, under small *B*_T_ and high 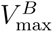 (Fig. S2).

**Figure 3:**
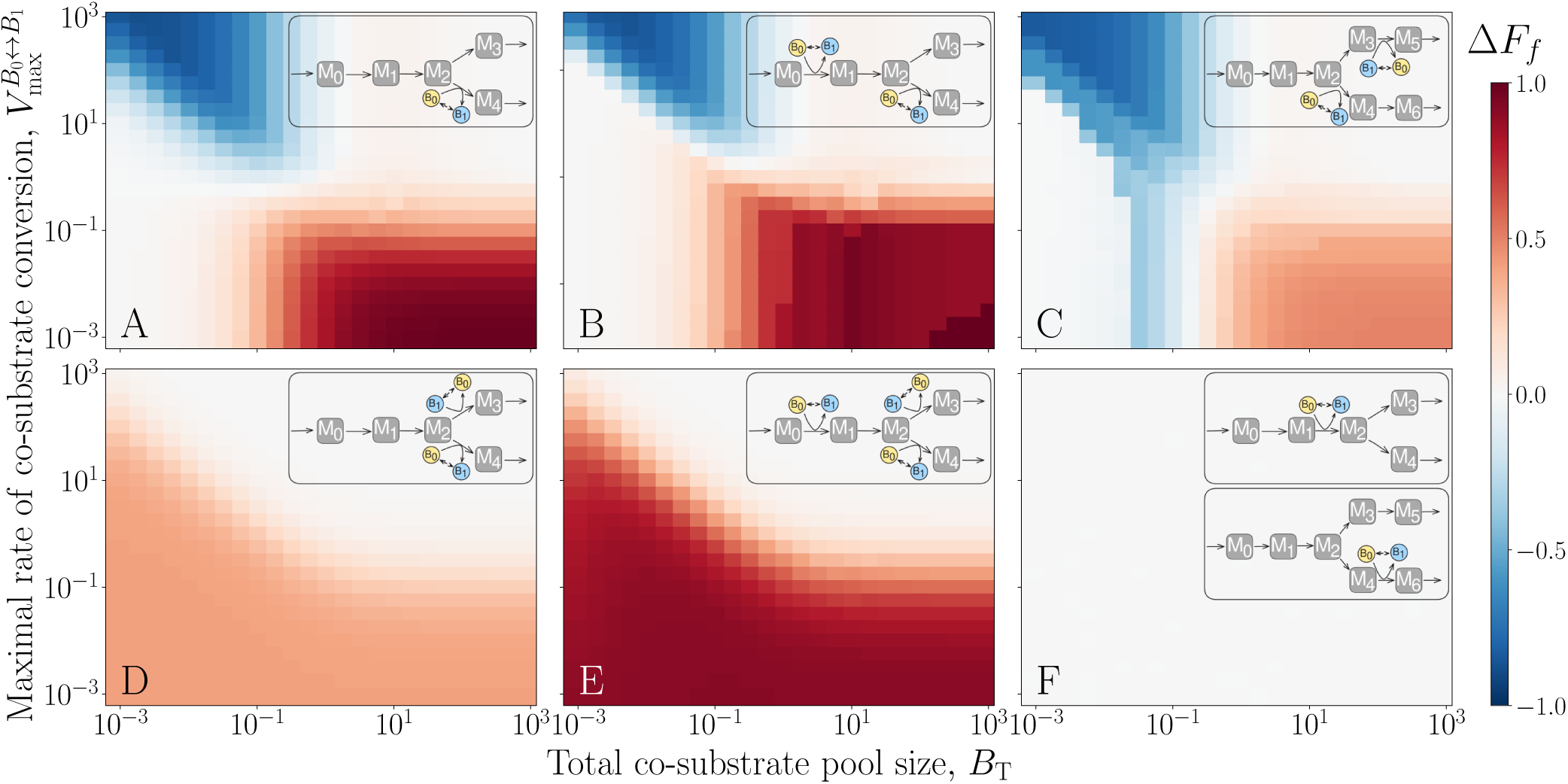
Motif structure influences capacity for regulation. **(A-F) Heatmaps showing** Δ*F_f_* = *F_f_* (max(*v*_in_))*−F_f_*(min(*v*_in_)) (defined in Fig 2B(ii)) as a function of *B*_T_ and 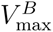 for different branch-point motifs: with a single CR at the branch point (similar to Fig.2C) (A); two CRs, one at the branch point and one upstream, with both using the same form of the co-substrate (B); two CRs, one CR at the branch point and the second downstream on the other branch, using complementary forms of the co-substrate (C); two CRs, one on each branch at the branch point and using complementary forms of the co-substrate (D); three CRs, two complementary CRs at the branch point and one upstream (E); and single CR either downstream of the branch point or upstream only (F). For each panel, Δ*F_f_* values are obtained by simulating the given system multiple times, for each combination of *B*_T_ and 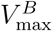, and against increasing *v*_in_ (as done in Fig.2). All parameters were set to 1, except for 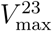 = 0.1. Notice that this parameterization creates a baseline bias in branch-point flux distribution favoring the bottom branch. Removing this bias and repeating the analyses gives qualitatively similar results (Figure S4).

### Flux switch requires asymmetric co-substrate usage at the branch point and is affected by down/upstream CRs

We next quantified the switch response Δ*F_f_* in different branch-point motifs, as a function of co-substrate pool size *B*_T_ and turnover rate 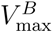. We find that high Δ*F_f_* can be achieved as long as at least one CR is placed at a branch point (Fig. 3A-E, see also Fig. S3 and S4). If a CR is placed anywhere other than immediately after the branch-point, then no switch response is produced as there is no asymmetric substrate usage at the branch point (Fig. 3F and S5). We find that Δ*F_f_* response is altered by additional CRs depending on how they alter the existing co-substrate usage. If the additional CRs exacerbate the co-substrate limitation at the branch point, then the response becomes more drastic (broader dark red region in Fig. 3B compared to A due to an additional upstream CR further depleting co-substrate *B*_0_). In contrast, if the additional CRs makes the co-substrate more available, i.e the two CRs now facilitate each other in terms of co-substrate availability, then the response magnitude Δ*F_f_* decreases (Fig. 3, compare A with C). When CRs that facilitate each other are both at the branch point, additionally the negative Δ*F_f_* region is lost (compare Fig. 3A to D). Adding an upstream CR into this motif re-enhances the asymmetry and once again increases the response (Fig. 3 compare D and E). We conclude that asymmetric co-substrate usage at the branch point is the key to the CR-driven flux switch (Fig. 3). This conclusion extends to other more complex motifs, in a way that can be explained by how the additional CRs affect the availability of co-substrate at the branch point CR (Fig. S6). Thus, a branch point CR provides a flexible mechanism for inducing nonlinear flux switch with increasing influx, and this switch response can be understood from asymmetric co-substrate usage.

### A simplified model of the yeast pyruvate branch point with co-substrate dynamics captures metabolic overflow and its experimental manipulation

The motifs analyzed above are similar to those seen in central carbon metabolism (see Fig. 1A-B). In the case of glycolysis, NAD^+^ reduction at the pyruvate branch point initiates the respiratory pathway, while in the fermentation pathway, NADH is oxidized further downstream (Fig. 1A). Upstream of pyruvate is another reaction utilizing NAD^+^. Thus, we turn into yeast to examine if and how NADH dynamics can induce flux switching (i.e. high Δ*F_f_*) with increasing influx, as experimentally observed in the metabolic overflow phenotype of this species (Fig. 4C, see also *Methods*).

**Figure 4:**
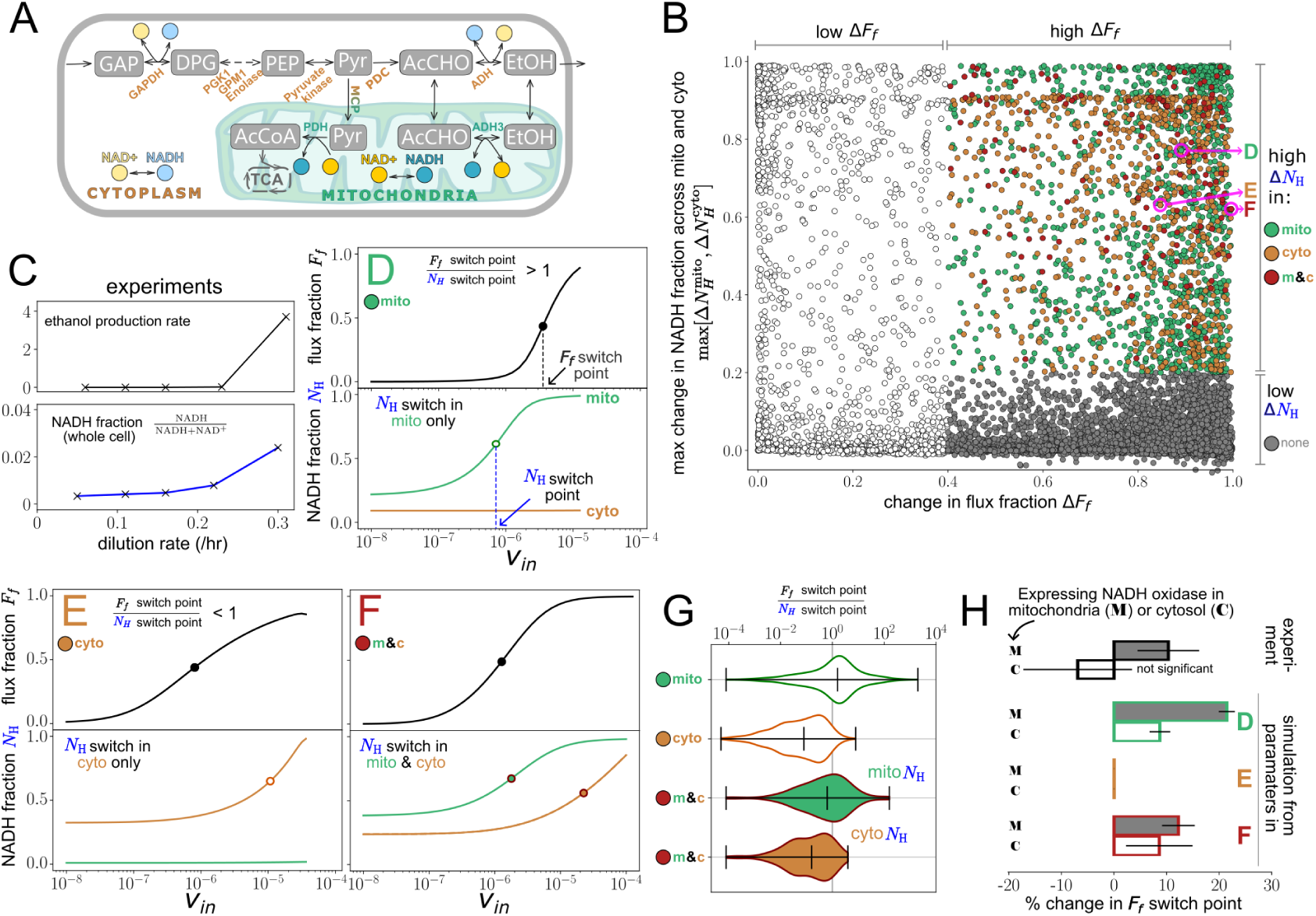
Yeast compartmental model. **(A)** A cartoon of the pyruvate branch point in yeast. Specific reactions are labelled with the relevant enzyme (orange and green texts). Dashed reaction line indicates multiple reactions have been condensed into one in the model. **(B)** Scatter plot of Δ*F_f_* vs. Δ*N*_H_ (the maximum among the mitochondrial or cytosolic [NADH]/([NAD^+^]+[NADH])). Each point shows results obtained when the model is simulated with a different parameter set, drawn from the ranges shown in Table S1. Individual simulations are classified as displaying: no switching (white), or switch in *F_f_* accompanied by a switch in mitochondrial NADH (green), cytosolic NADH (orange), or both (red). Magenta circles highlight examples cases from each category, analysed in (D-F). **(C)** Experimental data on the ethanol production rate^5^ and whole-cell NADH fraction^19^ against increasing growth rate (dilution rate). **(D-F)** The steady-state dynamics against increasing *v*_in_ for the simulation/parameter cases highlighted in magenta in (B). Each panel shows the plot of *v*_in_ vs flux fraction (top, black line) and co-substrate fractions (bottom, green and orange lines for mitochondrial and cytosolic NADH fraction, respectively). The switching points for *F_f_* and NADH fraction are marked with a circle on each panel, and on panel (D) the corresponding switch-point *v*_in_ values are marked. **(G)** Violin plots showing the ratio of the switch-point *v*_in_ for *F_f_* vs. that for NADH. Data is collated from different classes of simulations that show *F_f_* switch and: mitochondrial (green outline) or cytosolic NADH switch (orange outline), or both. The last case is analyzed using mitochondrial (green filled) or cytosolic (orange filled) NADH switch threshold. **(H)** Box plots showing the % change in threshold influx switch point in experiments (black outlines, data from^33^) or in the model (colored outlines). The upper and lower bars in each set show the effect of expressing NADH oxidase in the mitochondria vs. cytosol. Model results are collated from different classes of simulations that show *F_f_* switch and: mitochondrial (D, green outline) or cytosolic (E, orange outline) NADH switch, or both (F, red outline). For each class, simulations were performed with NADH oxidase expression mimicked in the mitochondria or cytosol by reducing 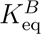 in the corresponding compartment (i.e. biasing the background reaction towards production of NAD^+^).

We created a simplified model of the pyruvate branch point in yeast (Fig. 4A). This model takes into account the fact that the pyruvate branch point is split across the cytosol and mitochondria, where pyruvate is transferred into mitochondria (via mitochondrial pyruvate transporter, MCP) before entering the TCA cycle, and crucially, the NAD^+^/NADH pools for the cytosol and the mitochondria are separate but connected by mitochondrial shuttles^38^. The model also accounts for the observed allosteric regulation (substrate activation) of PDC, located at the pyruvate branch point^24^.

We used parameters determined *in vitro* and utilized in previous models of yeast glycolysis^39^ where possible (see *Methods* and Table S1). We note that between the two branch enzymes, PDC has a higher *V*_max_ (high speed) and higher *K*_M_ (lower substrate affinity) compared to MCP, and that PDC is allosterically regulated by pyruvate, suggesting that mechanisms of enzyme asymmetry and allostery noted within the above didactic models (Fig. 2) should be relevant here. For parameters that are unknown, we sampled across physiologically relevant ranges^40,41^. For each set of sampled parameters, we quantified flux-switch extent, Δ*F_f_* after simulating the *F_f_* response against increasing influx. Inspired by experimental observations of changes in NADH across overflow initiation (Fig. 4C, bottom panel), we also quantified any switch-like dynamics in the NADH fraction (*N_H_* switch) by measuring the NADH fraction (i.e. NADH/(NADH + NAD^+^)) over the same influx range.

The model displays flux switching (high Δ*F_f_*) for many, but not all, parameter sets. Since the experimental value of Δ*F_f_* is unknown, we have arbitrarily defined cases where Δ*F_f_ >* 0.4 as displaying a flux switch. Among these, we further focus on the subset where high Δ*F_f_* is accompanied with an increase in the NADH fraction, i.e. an *N_H_* switch, occurring in the cytosol, mitochondria, or both (orange, green, and red dots in Fig. 4B). We focus on these subsets with an *N_H_* switch because in yeast chemostat experiments (Fig.4C), an abrupt increase in whole-cell NADH fraction accompanied, and possibly even preceded, the onset of metabolic overflow). Examples of the cases with different *F_f_* and *N_H_* switching dynamics are shown in Fig.4D-F. We define the ”flux switch point” as the influx value at which half-maximal flux change is realized, as illustrated in Fig.4D.

Strikingly, when mitochondrial NADH undergoes a switch (Fig.4D green), the *N_H_* switch point occurs at a lower influx than the *F_f_* switch point, suggesting that mitochondrial NADH dynamics may drive, rather than merely follow, the flux switch. In contrast, the cytosolic *N_H_* switch always occurs at a higher influx than the *F_f_* switch point (Fig.4E,F, compare orange with black), suggesting that cytosolic NADH dynamics do not drive flux switching. To further explore this difference in dynamics, we quantified the distribution of the ratio of the *F_f_* switch point to the *N_H_* switch point, for simulations where both switches happen (Fig.4G). This ratio exceeds one frequently in the cases displaying mitochondrial *N_H_* switching, whereas it is generally less than one in the cases with only a cytosolic *N_H_* switch (Fig.4G). Thus, the NADH switch might be driving the flux switch in the cases displaying mitochondrial *N_H_* switching. The associated parameters for the cases showing a mitochondrial NADH switch differ from the other cases mostly in their mitochondrial NADH turnover, having a lower turnover rate and higher K*_m_* for NAD^+^ (Fig.S7), which aligns with the expectation that the switch dynamics are due to mitochondrial NADH dynamics in these cases.

Given the effect of parameters controlling NADH turnover - both in this model and in the didactic models - we analyzed specifically the effect of altering NADH turnover on the flux switch point. For the cases displaying mitochondrial *N_H_* switch, increasing NADH oxidation in mitochondria (essentially deterring *N_H_* switch) pushes flux switch point to a higher influx (Fig. 4H, upper green outlined bar). For the same cases, when we increased NADH oxidation in the cytosol, we observed a similar shift in *F_f_* switch point, but this shift was smaller (Fig.4H, green outlined bars, lower bar). For the cases where only cytosolic NADH switching were observed, alterations of NADH turnover - either in mitochondria or the cytosol - had no effect (Fig. 4H orange outlined bars). These results match experimental findings in the budding yeast, where introducing a foreign NADH-oxidation reaction in the mitochondria shifts the metabolic overflow initiation point to higher glucose levels, but no significant effects were observed from introducing this reaction to the cytosol^31,33^ (Fig. 4H, black outlined bars labeled ”experiment”). In the model, we can also analyze the effect of increasing NAD^+^ consumption. Doing so in the mitochondria predicts shifting of the metabolic overflow initiation point to lower glucose levels (Fig. S9).

Finally, we wanted to further disentangle the contributions from differential enzyme kinetics and allosteric regulation of PDC versus contributions from NADH dynamics in driving switch dynamics. To this end, we considered models with or without co-substrates and with or without allostery/kinetic differences. In the absence of allostery/kinetic differences, we found instances of high Δ*F_f_* only with co-substrates (Fig.S8). In the presence of allostery/kinetic differences, high Δ*F_f_* could still be achieved without co-substrates, however, these models could not capture the observed changes in NADH during metabolic overflow or the effect of NADH oxidases observed in experiments.

Overall, these qualitative agreements between the model behavior and several experimental observations in yeast indicate that asymmetric dynamics arising from NADH-NAD^+^ usage at the pyruvate branch point and inside the mitochondria can drive the metabolic overflow dynamics in yeast, working separate to, but alongside, differential enzyme kinetics and allosteric regulation of PDC.

## DISCUSSION

Here, we analyzed the flux dynamics at a metabolic branch point, with or without co-substrate-utilizing reactions at and around the branch point. Using mathematical models, we have shown three independent mechanisms enabling flux switching at the branch point with increasing influx: (i) differential kinetic parameters for the branch-point enzymes, (ii) allosteric regulation of one of the branch-point enzymes, and (iii) presence of co-substrate-utilizing reactions at and around the branch point. These mechanisms constitute a type of self-regulation, as they do not require active control of enzyme levels and arise solely from pathway structure and placement of co-substrate reactions. To see how these mechanism could apply to overflow metabolism seen in yeast, we developed a specific model of the pyruvate branch point in this species, and the NADH co-substrate usage around this branch. Using measured or physiologically feasible kinetic parameters, we found a flux-switching response with increasing influx through all three mechanisms to be possible in this model. However, only the co-substrate-based mechanism is capable of explaining existing experimental data from yeast on NADH and metabolic overflow dynamics, and their influence on each other. Taken together, these findings demonstrate that branch points and co-substrate utilization around them act as key, ’built-in’ regulatory elements within cellular metabolism.

The qualitative agreement between the presented model and yeast metabolic overflow dynamics provides strong support that co-substrate dynamics could influence metabolic overflow. The observed increase in NADH levels in yeast during metabolic overflow is captured by the co-substrate model. It must be noted, however, that the experimental measurements pertain to the whole cell, rather than mitochondria and cytosol separately, while the model indicates a larger impact of mitochondrial NADH dynamics to metabolic overflow initiation (Fig. 4(B)). Testing the effects of NADH dynamics on the threshold for metabolic overflow, the experiments revealed that only mitochondrial NADH oxidases can shift the metabolic overflow threshold. The cited study mentions expressing cytosolic NADH oxidase did not yield significant results and error bars are accordingly large in Fig. 4H). These experimental findings, as well, agree with the presented model (Fig. 4H and S9.

The explanatory power of the presented theory, based on NADH dynamics, can extend to other observations on metabolic overflow in yeast and other species. In this theory, the metabolic overflow initiation point is predicted to tightly depend on NADH-NAD^+^ turnover rates and pool sizes. Thus, these key physiological parameters in different species or strains, shaped by adaptations to past environmental conditions such as stable or changing oxygen levels^15^, can determine their metabolic overflow phenotypes. In this context, we note that recent proteomics studies showed that Crabtree negative yeasts, which have a higher threshold for metabolic overflow, display significantly higher expression levels of electron transport chain enzymes^15,42^. In light of the presented models, this observation can be interpreted as higher mitochondrial activity allowing for faster NADH turnover (into NAD^+^) enabling higher tolerance of glycolytic influx at the pyruvate branch point. Indeed, the NAD^+^/NADH ratio was found to be higher in Crabtree negative vs. positive yeast^15^. The presented theoretical findings also align with the findings from bacteria, where altering of NADH dynamics resulted in shifting of the metabolic overflow initiation point in *Escherichia coli* ^33,34^. In mammalian cells as well, alterations in NADH dynamics are found to affect flux distributions both at the pyruvate branch point, influencing overflow metabolism^25^, and downstream at the alpha-ketoglutarate branch point, influencing the induction of reductive glutamine consumption^35^. These diverse finding from different species, all linking to NADH dynamics support the broader notion of co-substrate dynamics acting as flux regulators across branch points.

The presented theory provides a clear mechanistic basis on how co-substrate dynamics can affect metabolic flux distributions. As such, it is linked to a series of historic studies proposing a regulatory role for co-substrates^43,44^, including the phenomenological notion of ATP acting as an indicator of cellular ‘energy charge’^44–46^. More recent studies have shown that co-substrate dynamics can act as flux limiting^28–30^, adding to a body of literature showing that co-substrate dynamics give rise to nonlinear dynamics within metabolism . The presented work extends these studies by showing a direct nonlinear regulatory effect of co-substrate dynamics around branch points. While we considered here specifically the pyruvate branch point, and NADH dynamics, it is noticeable that key junctions in central metabolism such as the folate cycle, underpinning single carbon transfers, and the TCA cycle, underpinning nitrogen assimilation, involve branch points with co-substrate usage around them. Thus, future studies can expand the modelling of co-substrate dynamics to specific branch points and also to capturing the interlinked nature of multiple branch points and multiple co-substrates acting on them. There is already experimental findings suggesting, for example, that aerobic glycolysis in mammalian cells involves an interplay between NAD^+^-NADH and ATP-ADP dynamics^25^.

The dynamical explanation for metabolic overflow we presented here is distinct from those explaining metabolic overflow arising from cells altering their protein investment into fermentative and respiratory pathways, according to the proteome efficiency of these two pathwaysThose theories operate at a higher, phenomenological level, and do not provide any mechanistic explanations for how cells sense and initiate such a proteome allocation adjustment and how different species or strains can have different metabolic overflow thresholds^1^. Thus, the presented theory could be complementary to the ideas surrounding protein allocation, as co-substrate-based flux regulation at branch points can act as a fast, self-regulatory response to achieve flux changes, which can then be facilitated further with slow, and longer-term, changes in enzyme allocation. This idea aligns with experimental findings on metabolic overflow in yeast, whereby time-resolved perturbation of dilution rate in a chemostat resulted in a metabolic overflow within minutes and transcriptional changes are only observed in hours, after adaptation to increased dilution rate^4^. It would, therefore, be interesting to expand ongoing efforts of generating ‘coarse-grained’ cellular models, based on resource allocation, with the inclusion of co-substrate dynamics and key branch points from central metabolism.

In recent years, attempts are emerging to modulate co-substrate dynamics for direct engineering of cellular metabolism for biotechnological applications^47–49^. We note that the presented co-substrate-based theory - and extensions of it - can provide a predictive basis for these attempts. In particular, specific modulation of NADH turnover rates can be used to engineer metabolic overflow through the pyruvate branch point. Future application of the presented theory to other branch points, or several connected branch-points, can allow engineering of metabolic fluxes associated with other phenotypes.

## METHODS

### Analysis of branch-point motifs in genome-scale models

We used latest genome-scale metabolic models for yeast^36^ and *E. coli* ^37^. For a given model, we identified those metabolites that are consumed in two or more reactions, excluding excretion reactions but including inter-organelle transport reactions. These metabolites are counted as ”branch-point metabolites”. We then categorized each branch-point metabolite as involving a CR, if any of its immediate consuming reactions involved co-substrates AT(D)P or NAD(P)H (and their different forms). The analysis was performed for all compartments, but the results are shown only for the cytosol as this group accounted for most of the cases identified. The analysis script, written in MATLAB, as well as the genome-scale models used are available at the Github repository: https://github.com/OSS-Lab/Co-substrate-induced-flux-switching-at-branch-points-can-explain-overflow-metabolism

### Modelling of branch-point motifs

Our modelling approach is summarized visually and mathematically in Fig.S1. In brief, we used ordinary differential equations (ODEs) to model the dynamics of branch point containing pathways, with and without CRs and with and without allostery involving the branch point metabolite. Main pathway reactions are modeled using single or two-substrate versions of the Michealis-Menten rate equation (see Fig.S1).

In the case of CRs, specific co-substrate forms - generically represented as *B*_0_ and *B*_1_ - are assumed to take part in a given reaction and the total amount of co-substrate is assumed to be fixed (i.e. *B*_0_ + *B*_1_ = *B*_T_). Relaxing this assumption does not alter qualitative nature of the conclusions presented here (see Fig.S10). In addition to the CRs directly modeled as part of pathway motifs, a ’background’, reversible *B*_0_↔*B*_1_ conversion reaction is implemented to account for reactions not related to the branch-point motif, with an equilibrium coefficient of 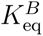 .

This reaction is modeled using the reversible Haldane rate equation (see Fig.S1). The resulting reactions and rate equations are incorporated in a series of ODEs for each model and depending on the model architecture. A set of ODEs for an example system are given in Fig.S1. All ODE systems are implemented as a Python script and solved using numerical methods implemented in scipy library. All model and analyzes scripts are made available at the following Github repository: https://github.com/OSS-Lab/Co-substrate-induced-flux-switching-at-branch-points-can-explain-overflow-metabolism

### Simulation of flux behavior

A key metabolic behaviour we have focused on here is the dynamics of flux distribution at a branch point. To quantify this behavior, we considered the steady-state flux fraction into the upper branch as a key measure describing the pathway dynamics, (*F_f_* = *F*_upper_/(*F*_lower_+*F*_upper_)). To get *F_f_* for a given pathway architecture, we simulate the ODEs to steady state for increasing values of influx rate, *v*_in_, and record system variables, resulting in a response curve as shown in Fig. 2. Simulated systems are assumed to have reached steady state, when all metabolite gradients are smaller than 10*^−^*^5^ after a simulation time of 10^5^ minutes. It is shown previously that attainment of steady-state in a metabolic pathway with co-substrates requires *v*_in_ to be lower than the maximal rate of enzymes in the pathway and a nonlinear function relating to *B*_T_ and 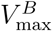 ^28,30^. Here, for different branch-point motifs and key parameter choices, we analyse *F_f_* within the steady-state regime, i.e. before *v*_in_ becomes larger than either one of these thresholds. For the models used in motif analyses, reaction kinetic parameters are set to arbitrary values for simplicity, with the key parameters, such as 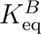, *B*_T_, and maximal rate of the ’background’ *B*_0_↔*B*_1_ conversion reaction varied to assess their effects in system dynamics (as shown in Fig. 2 and Fig.3). These phase plane analyses involve running time-course simulations as described above for different parameters sets, to steady state, and obtaining *F_f_* .

### Yeast pyruvate branch-point model

The simplified model of the lower glycolysis and the pyruvate-branch point in yeast are modeled based on previous models of the lower glycolysis in the same organism^32,39^. The model is simplified by only focusing on the pyruvate branch point, lumping some of the linear parts of the lower glycolysis into a single reaction, and expanding with reactions around the branch point. The model includes the compartmentalization of NADH across cytosol and mitochondria, resulting in distinct cytosolic and mitochondrial pools, along with NADH-shuttle reactions that are proposed to consume/produce NADH using either of these pools^38^. Transport of pyruvate into mitochondria is modeled and forms, together with the pyruvate decarboxylase mediated reaction in the cytosol, the pyruvate branch point in this model. Kinetic parameters for most reactions are obtained from the previous yeast model^39^. The kinetics of the pyruvate transport dynamics are obtained from measurements with isolated mitochondria^50^, while remaining kinetic parameters are obtained from databases^51,52^. To account for unknown parameters, and allow for variances in experimental measurements, many of the kinetic parameters are further sampled from a wide, physiologically relevant, range, and centered around measured values where available (see Table S1). For the background reaction for NADH turnover, the 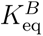 is set to achieve a NAD^+^ to NADH ratio of 100, as observed for the yeast cytosol, but also varied as low as 7, as observed for whole cell metabolite measurements^53^. When comparing the model with/without NADH oxidase, the switch point comparison was done in a way to account for any changes in Δ*F_f_* that might be caused by introducing NADH oxidase.

### Re-analysis of experimental data

Experimental data plotted in Fig 4C was obtained from two consecutive studies that measured metabolite levels across chemostat experiments, including during metabolic overflow initiation^5,19^. The two studies analyzed the same experimental samples^19^. Data points for ethanol production rate were extracted from the graph in Supplementary Figure S6 of^5^ using plotdigitizer. Absolute concentrations of NADH and NAD^+^ from LC-MS/MS analysis were extracted from Supplementary Dataset 9 in^19^. As per the published methods, each point indicates concentrations averaged over 4 technical replicates at any given dilution rate of a glucose-limited chemostat. As cell growth is limited by the dilution rate in a chemostat reactor, we use dilution rate under glucose limitation as a proxy for metabolite influx rate into glycolysis. Experimental data plotted in Fig 4H (upper panel) was calculated using switch-point dilution rates mentioned in the main text of^33^.

Cytosolic perturbation corresponds to the strain AOX, while mitochondrial perturbation corresponds to the strain NOX, described in^33^. The percentage change for each perturbation was calculated using the switch-point dilution rate of a control strain, described in^33^, as reference value, and error propagation was used to calculate the error bars for each perturbation.

## ACKNOWLEDGMENTS

This work was funded by the UK Biotechnology and Biological Sciences Research Council via grant BB/T010150/1, the Gordon and Betty Moore Foundation via grant GBMF9200, and the Leverhulme Trust grant id RPG-2024-387. We thank Elad Noor for useful discussions on reaction kinetic parameters and for comments on an earlier version of this manuscript. The authors also thank all members of their research groups for their support.

## AUTHOR CONTRIBUTIONS

OSS and RW conceived the study and developed mathematical models. S, WS and TDSGR provided resources and analysed experimental data. RW, S, and WS created figures. All authors contributed to writing the manuscript.

## SUPPLEMENTARY INFORMATION

Figures S1-S10 and associated legends

Table S1. Parameter ranges used in the yeast model.

