## Supplementary Information for "Asymmetric co-substrate usage at a metabolic branch point as a minimal network motif that can drive overflow metabolism"

August 5, 2026

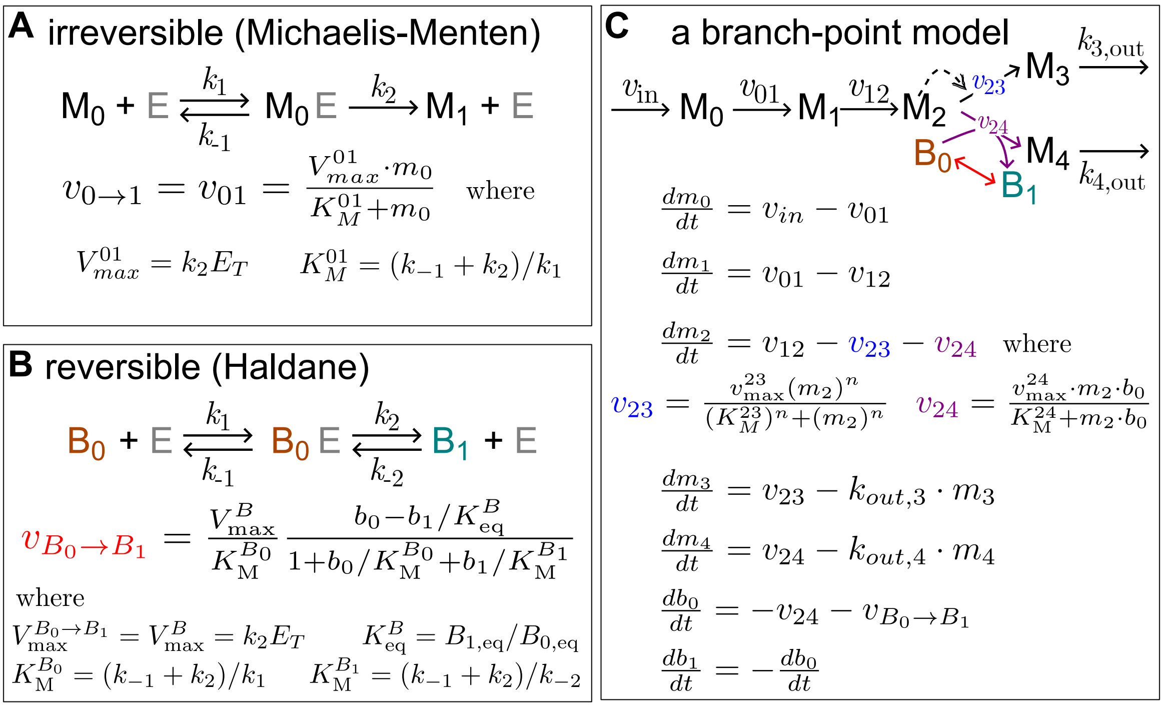


Figure S1: **Cartoon representation and details of the kinetic modelling of branch-point motifs. (A, B)** To model branch-point motifs, we consider simple pathway sections with influx (from upstream processes) at a rate $v_{\text{in}}$, and outflux of metabolites at a rate $k_{\text{out}}$. Reactions within the pathway section are modelled using the established Michealis-Menten rate equation, while background turnover of co-substrates, when present, are modelled using the reversible Haldane rate equation. $\text{M}, \text{B}$, and E represent metabolites, co-substrates and enzymes respectively; $k$ are binding/unbinding/reaction rates, $V_{\text{max}}$ is the maximal rate at large substrate concentrations, $K_{\text{M}}$ is the Michaelis-Menten constant (substrate concentration at which half maximal rate is achieved) and $K_{\text{eq}}$ is the equilibrium constant for the specified reaction (i.e. the ratio of product vs substrate when the reaction has reached steady state). Total co-substrate level ($B_0 + B_1$) is fixed. We find that this assumption can be relaxed, and changes in co-substrate total pool can be incorporated into the models without loss of generality (Figure S10). The effects of any 'off-pathway' reactions consuming and generating the co-substrate are modeled as a single, enzymatic reversible reaction from $B_0$ to $B_1$ with equilibrium coefficient $K_{\text{eq}}^B$ and maximal rate $V_{\text{max}}^B$. **(C)** Schematic and ODEs for the branch-point model showing a reaction with substrate-dependent cooperativity ($M_2$ to $M_3$, with $n > 1$) and a reaction with co-substrate ($M_2$ to $M_4$). In the latter case, we assume that the substrate, product and their associated co-substrate bind/unbind simultaneously, resulting in the reaction rate being the same as for single substrate reactions.

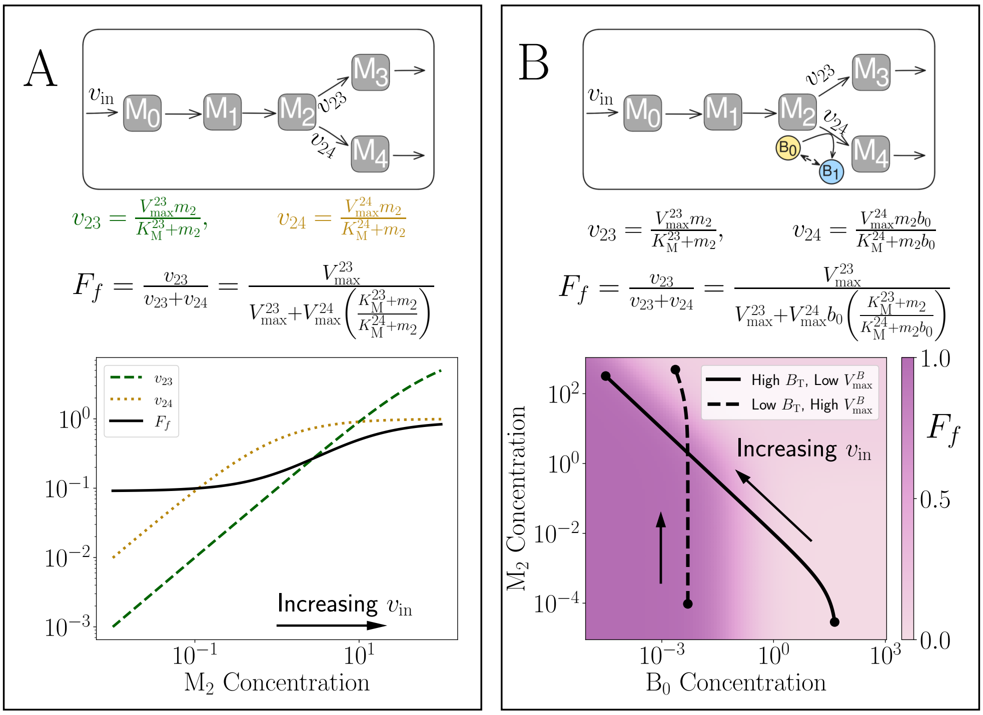


Figure S2: **Theoretical results for flux fraction dependence on $v_{in}$.** (A) Case without co-substrates. Line plots show the flux from $M_2$ to $M_3$ ($v_{23}$; green dashed line), $M_2$ to $M_4$ ($v_{24}$; yellow dotted line) and the flux fraction into $M_3$ ($F_f$; black line). Parameters are $(K_M^{23}, K_M^{24}, V_{max}^{23}, V_{max}^{24}) = (100, 1, 10, 1)$. (B) Case with co-substrate utilization on the lower branch. Parameters are $(K_M^{23}, K_M^{24}, V_{max}^{23}, V_{max}^{24}) = (1, 1, 0.1, 1)$ (the same in as in Fig. 3A). Dotted/solid lines use $(B_T, V_{max}^B) = (0.01, 100)/(100, 0.01)$, from the blue/red regions of Fig. 3A respectively. In (B), the steady-state values for $b_0$ and $m_2$ are calculated from simulations, and black lines show the value of $b_0, m_2$ and $F_f$ as $v_{in}$ increases.

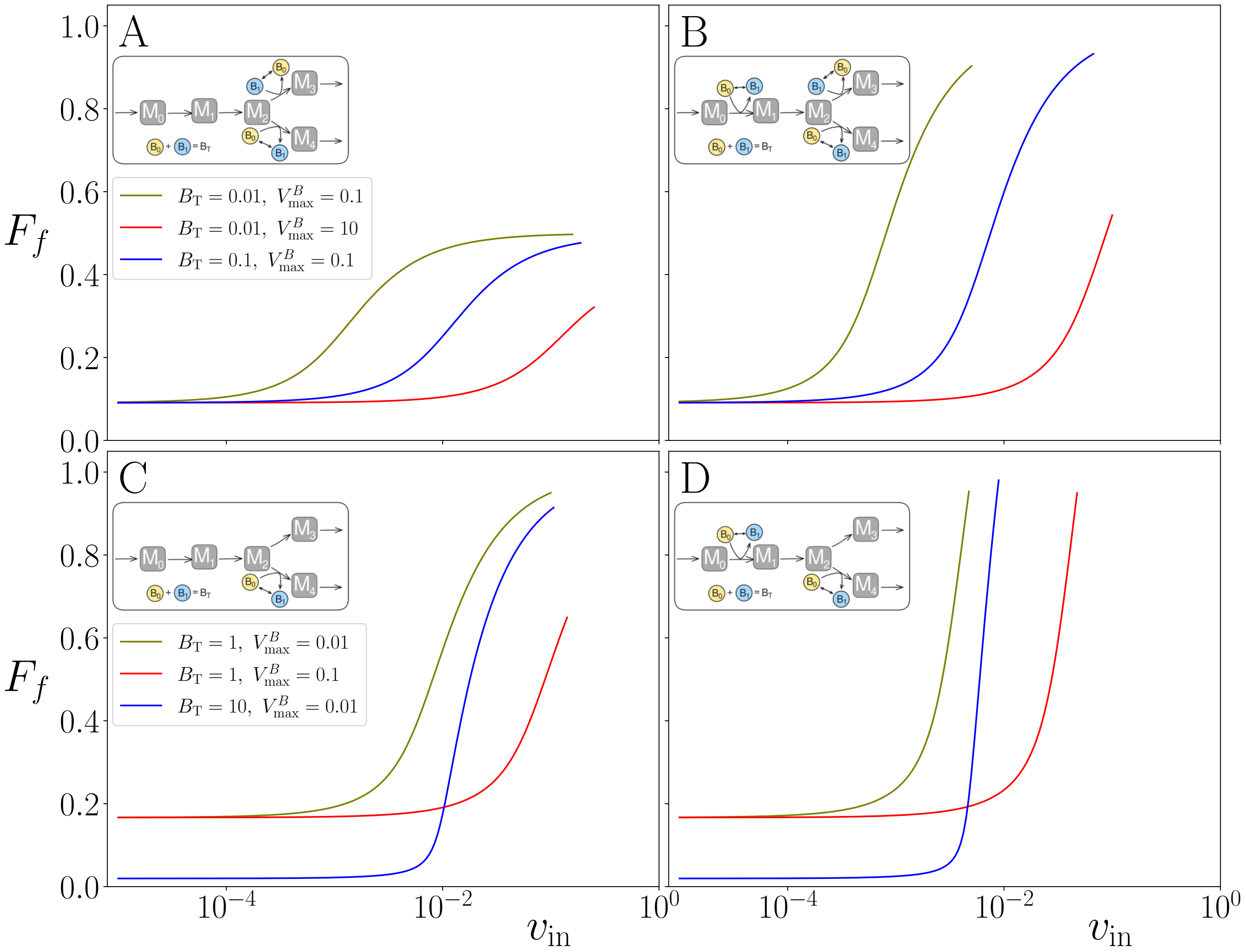


Figure S3: **Impact of co-substrate pool size and background $V_{\max}$on switching dynamics.** Flux fraction versus $v_{\mathrm{in}}$ for the motifs shown in the insets, some of which are also shown in Fig. 3. Simulation results in different colors are obtained with all parameters set to 1, except those parameters shown in the legend. Legend in panel A applies to both panels A and B, and the legend in C applies to both panels C and D.

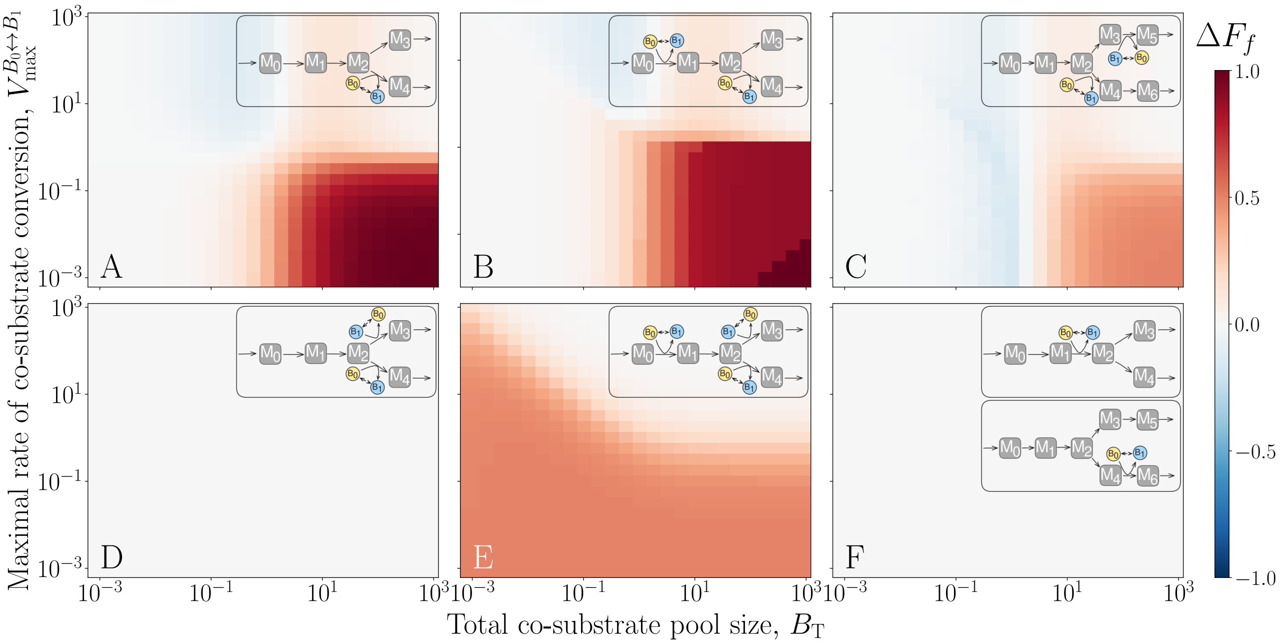


Figure S4: **Motif structure influences capacity for regulation even when branch enzymes have identical kinetics.** Heatmaps showing $\Delta F_f = F_f(\max(v_{\mathrm{in}})) - F_f(\min(v_{\mathrm{in}}))$ (as shown in Fig 2B(ii)) as a function of $B_{\mathrm{T}}$ and $V_{\max}^{B}$ for metabolic motifs with (A) a single CR placed on the branch point (similar to Fig. 2C); (B) two CRs, one at the branch point and one upstream, and both using the same form of the co-substrate; (C) two CRs, one CR directly at the branch point and the second downstream on the other branch, using the different forms of the co-substrate. (D) two CRs, one on each branch, using different forms of the co-substrate; (E) three CRs, two at the branch point and one upstream; (F) single CR either downstream of the branch point or upstream only. Simulations are performed for each combination of $B_{\mathrm{T}}$ and $V_{\max}^{B}$ with all other parameters set to 1 (i.e. no enzymatic bias towards either branch).

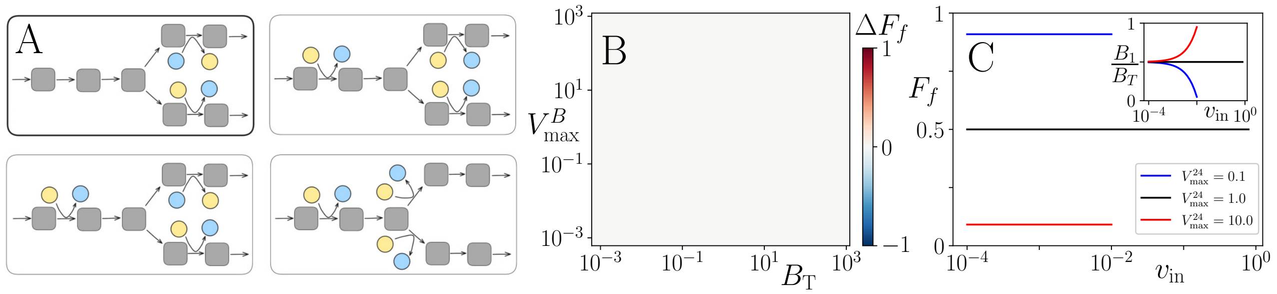


Figure S5: **Off-pathway or symmetric co-substrate usage does not result in regulation.** (A) Motifs showing placement of CRs around metabolic branch points that do not result in regulation as $v_{\mathrm{in}}$ varies (B) Heatmap of $\Delta F_f$ as a function of $B_{\mathrm{T}}$ and $V_{\max}^{B}$ for the motif with the bold outline in A. $(K_{\mathrm{eq}}^{B}, V_{\max}^{24}) = (0.1, 0.1)$, with all other parameters set to 1. (C) Flux fraction versus $v_{\mathrm{in}}$ for $(B_{\mathrm{T}}, V_{\max}^{B}) = (10, 0.01)$, $V_{\max}^{24}$ as indicated in the legend and all other parameters set to 1.

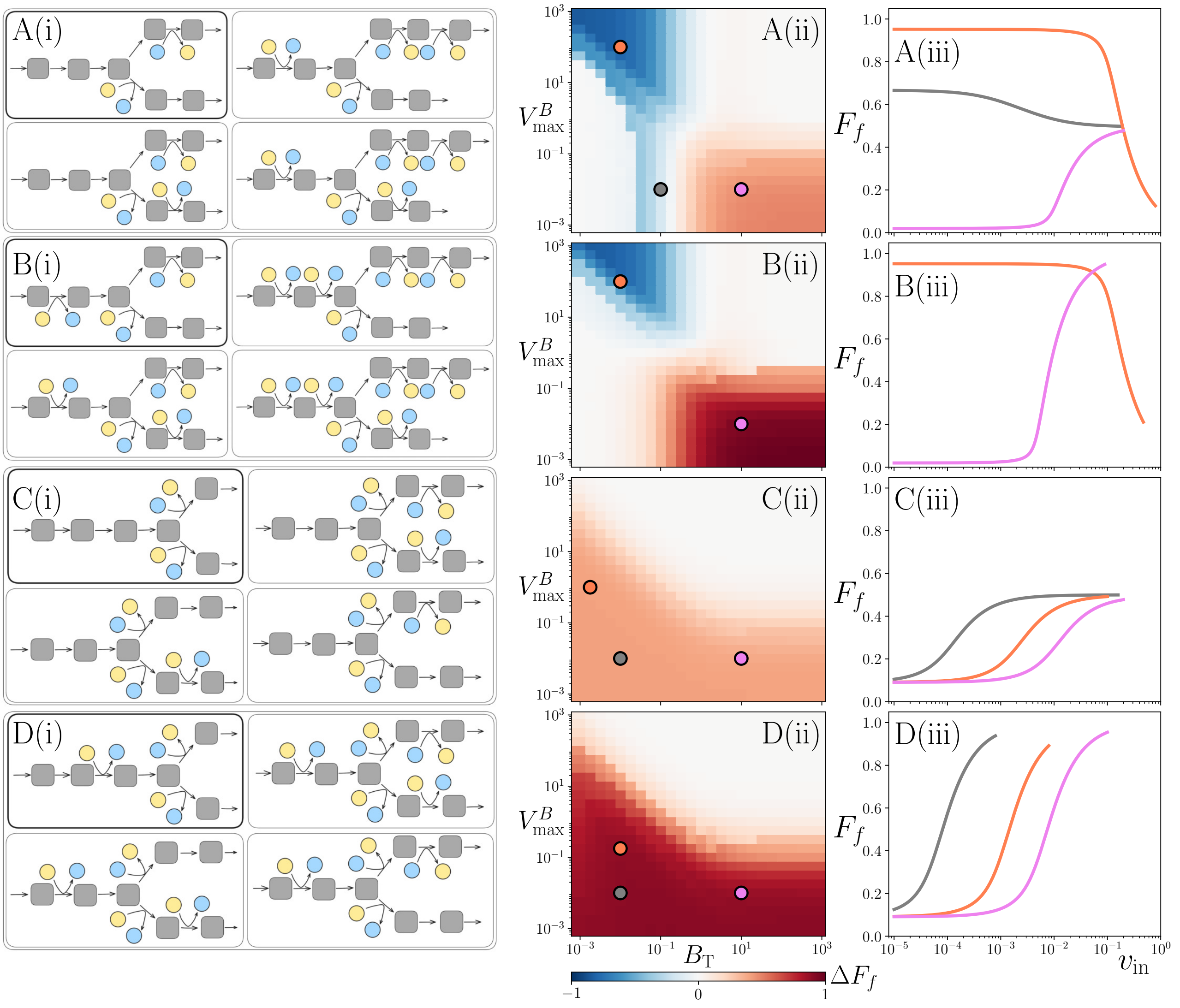


Figure S6: **Categorization of regulation types and associated motifs.** (A-D)(i) Motifs exhibiting similar regulation, as shown in the adjacent heatmaps. (A-D)(ii) Heatmaps showing $\Delta F_f = F_f(\min(v_{\text{in}})) - F_f(\max(v_{\text{in}}))$ for motif with bold outline in panel (i). All other motifs in each panel have similar heatmaps. $V_{\max}^{23} = 0.1$ (top branch $V_{\max}$) and all other parameters set to 1. (A-D)(iii): Flux fraction vs. $v_{\text{in}}$ for the bold motif in the same row, with parameters corresponding to the dots in the adjacent heatmaps.

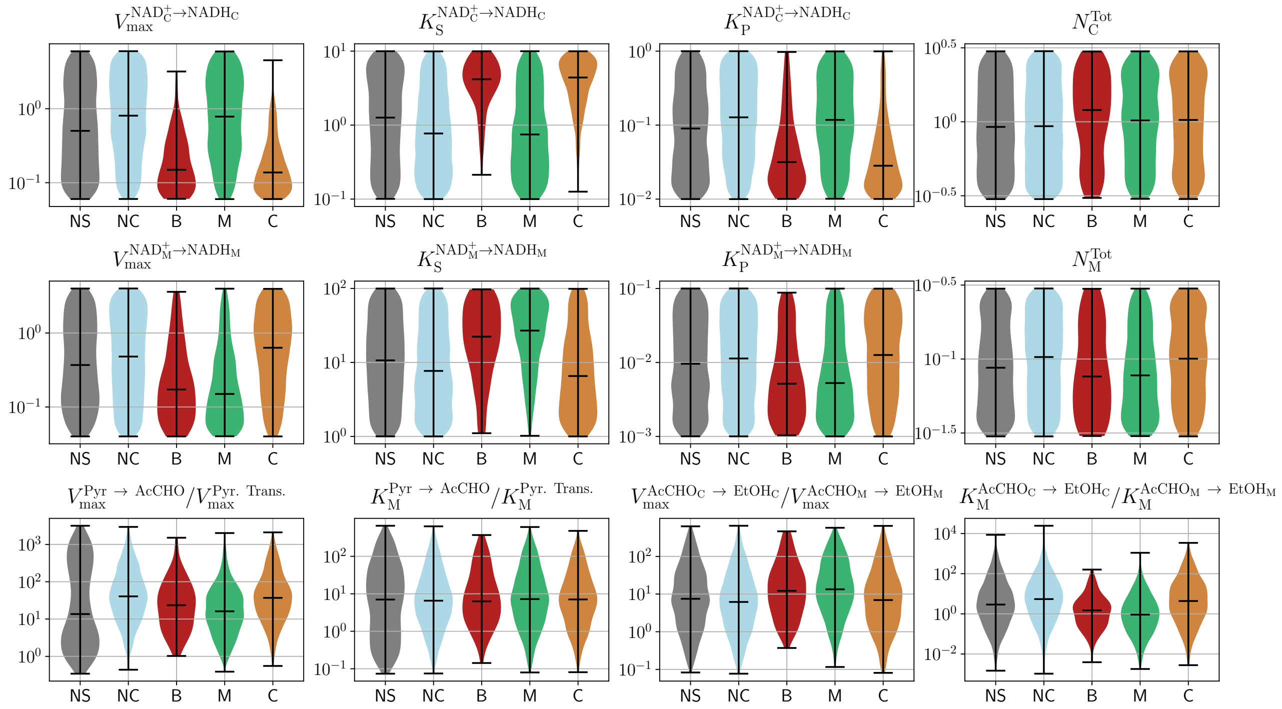


Figure S7: **Comparison of parameter regimes in the compartment model** Parameter distributions for the models with parameter sets displaying the 5 "switching types" identified in Figure 4: no switching (NS), no co-substrate switching (NC), switching in NADH fractions in both mitochondria ($N_{\rm H}^m$) and cytosol ($N_{\rm H}^c$) (B), switching in only $N_{\rm H}^m$ (M) and switching only in $N_{\rm H}^c$ (C). These sets are indicated on the x-axis of each panel and each panel shows the distribution for a different parameter as indicated on the panel title. Parameters are grouped as follows: top and middle row shows the distributions for the NADH background reaction parameters for cytosol and mitochondria, respectively, while the bottom row shows the ratios of $V_{max}$ and $K_m$ of the reactions at the pyruvate branch point and of mitochondrial vs. cytosolic ethanol reactions. $K_S$ and $K_P$ are Michaelis-Menten constants for the substrate and the product of a reaction, respectively.

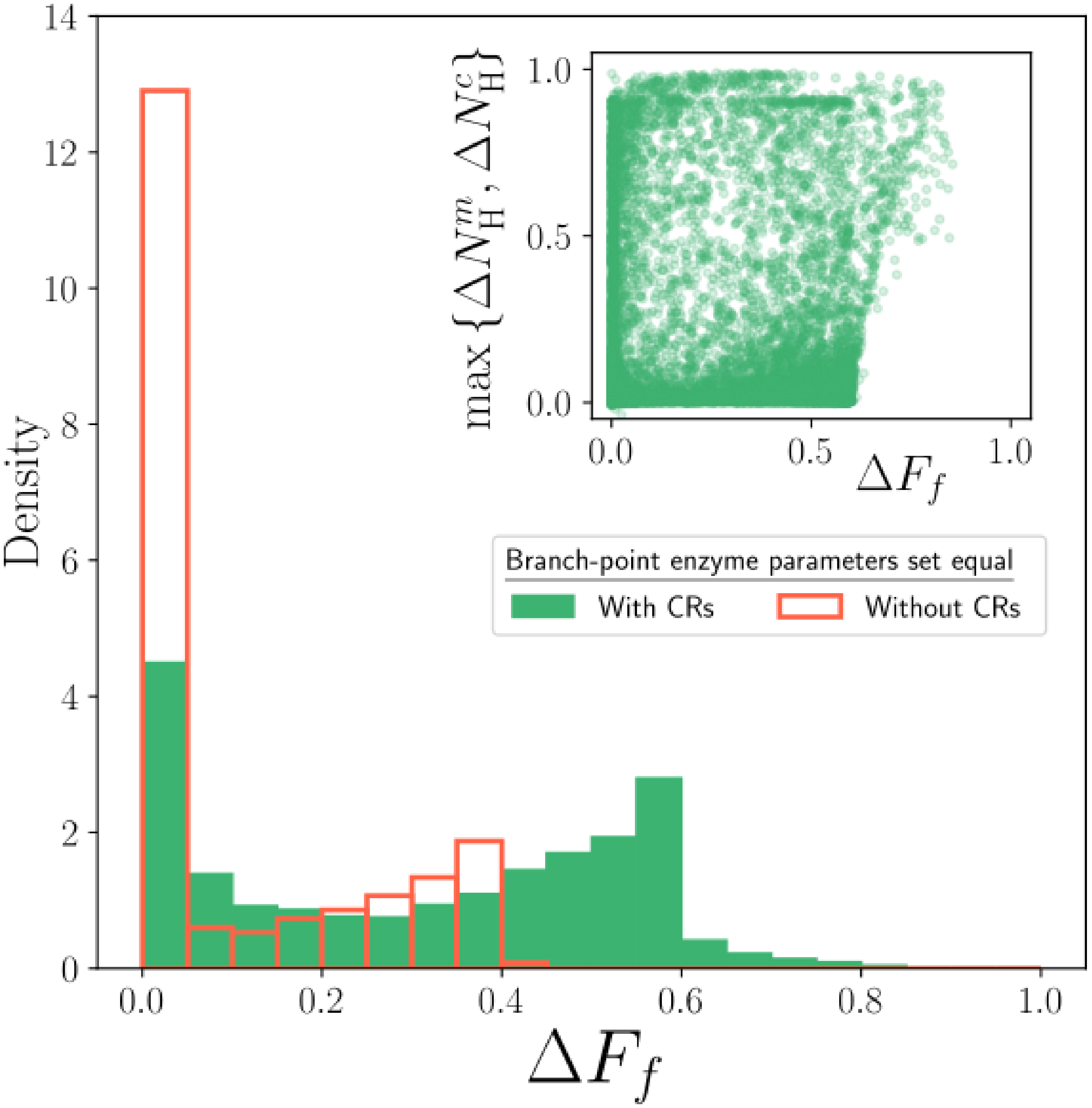


Figure S8: **Co-substrate dynamics alone can create switches in the compartment model** Histograms for the distribution of $\Delta F_f$ in the case with/without co-substrates (green filled/red outline) for the parameter regime shown in Table S1, but with all the reactions that have Pyruvate as a substrate set to have the same parameters to simulate the scenario without differential enzyme kinetics, and without allosteric activation of PDC by pyruvate.

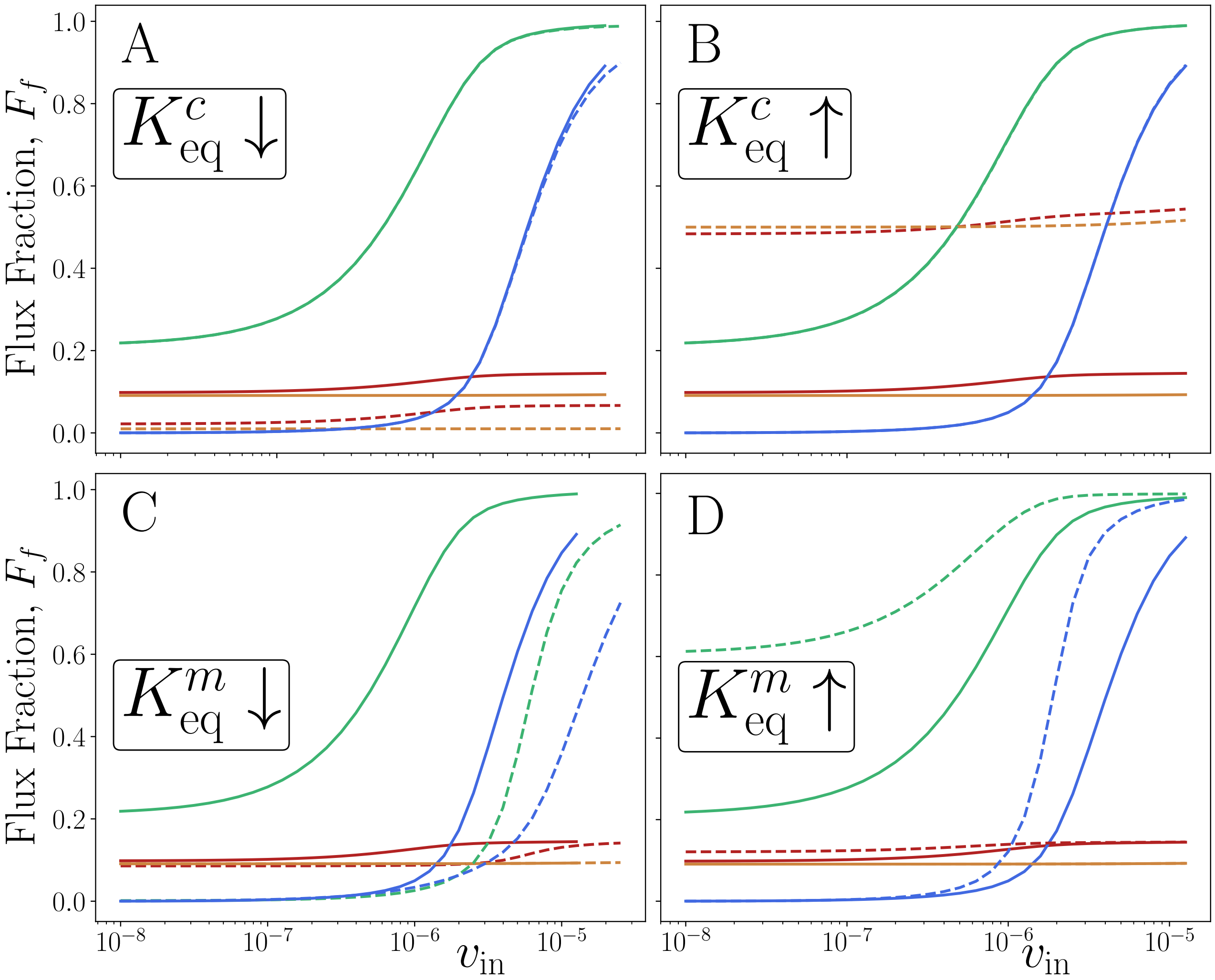


Figure S9: **Mitochondrial co-substrates drive switching** Effects of changing the NADH bias in the cytosol and mitochondria, in sample systems that are also shown in Figure 4. Solid lines are for the base model, without the $K_{\text{eq}}$ change, while the dashed lines show the effect of changing the cytosolic NADH levels (A,B) or the mitochondrial NADH levels (C,D). Blue lines show flux fraction, while green, orange, and red lines show the mitochondrial, cytosolic, and whole-cell NADH fractions, respectively. Altering NADH dynamics is modelled by changing the $K_{\text{eq}}$ for the background conversion (as indicated in each panel). Left column shows biasing the background reaction towards $\text{NAD}^+$, while the right column shows biasing towards NADH. Parameters are the same as in Fig. 4F, apart from the $K_{\text{eq}}$ as indicated in each panel. Increased/Decreased $K_{\text{eq}}$ means it is multiplied/divided by 10.

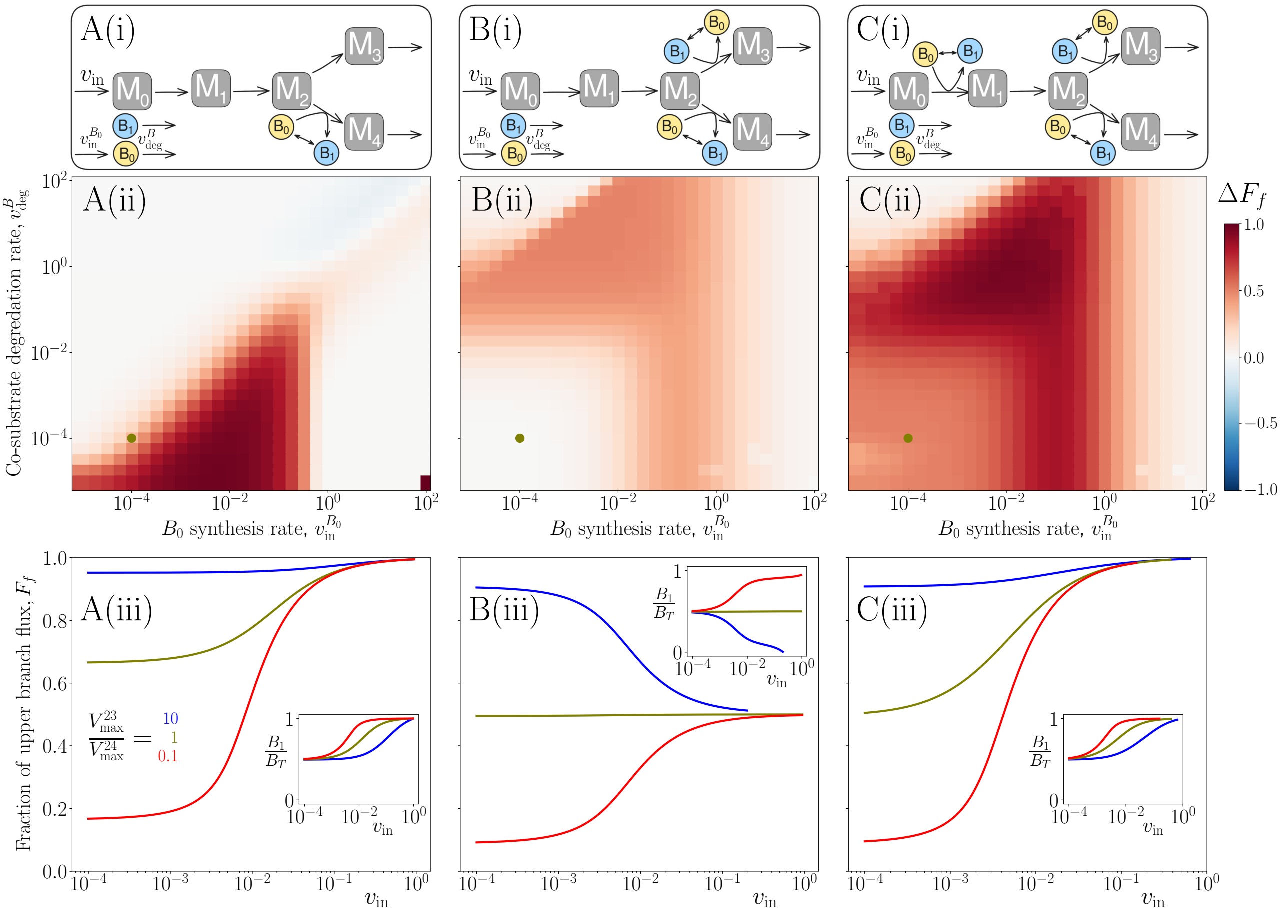


Figure S10: **Regulation persists in the presence of synthesis and degradation of co-substrates.** (A-C)(i): Simple branch-point motifs with co-substrates on only one branch (A), on both branches (B), and on both the branches and upstream (C). All reactions apart from the background conversion of $B_0$ and $B_1$ are modeled as irreversible. (A-C)(ii): Heatmaps showing $\Delta F_f = F_f(\min(v_{\text{in}})) - F_f(\max(v_{\text{in}}))$ as functions of $v_{\text{in}}^{B_0}$ and $v_{\text{deg}}^{B}$ for the metabolic systems in the panels above. $(B_{\text{T}}(t=0), V_{\max}^{B}) = (10, 0.01)$, and all other parameters are set arbitrarily to 1 (A-C)(iii): Simulation results from models corresponding to the motifs shown in A-D. Each panel shows the steady-state flux fraction from $M_2$ to $M_3$ versus the influx into the pathway $v_{\text{in}}$. Red,olive and blue lines indicate conditions favoring the lower branch, neither branch or the upper branch, respectively. Parameters are the same as in the panels above, with ($v_{\text{in}}^{B_0}$, $v_{\text{deg}}^{B}$) corresponding to the olive dot, Red/olive/blue lines indicate when the $v_{24}$/neither/$v_{23}$ reaction is favored and all other parameters are set arbitrarily to 1. Insets in (A-C)(iii) show the $B_1/B_{\text{T}}$ fraction, with the different colored lines showing simulation results with parameters as in the main panel.

| Michaelis-Menten (Reversible): $\nu = \frac{V_{\max}}{K_{\mathrm{M}}^{\mathrm{S}}} \frac{s - p/K_{\mathrm{eq}}}{1 + s/K_{\mathrm{M}}^{\mathrm{S}} + p/K_{\mathrm{M}}^{\mathrm{P}}}$ | | | | | | |
|---|---|---|---|---|---|---|
| Reaction (enzyme) | $V_{\max}$ ($\mu$mol/min/mg) | $K_{\mathrm{M}}^{\mathrm{S}}$ $(mM)$ | $K_{\mathrm{M}}^{\mathrm{P}}(mM)$ | $K_{\mathrm{eq}}$ | Pool size $(mM)$ | Citation(s) |
| GAP $\xleftrightarrow{\text{gapdh}}$ DPG | 1 – 10 | 0.001 – 1 | 0.001 – 1 | 0.0056 | | [1] |
| DPG $\xleftrightarrow{\text{pgk1, gpm1, enolase}}$ PEP | 1 – 10 | 0.001 – 1 | 0.001 – 1 | 0.1 – 10 | | [1] |
| PEP $\xleftrightarrow{\text{pyruvate-kinase}}$ PYR | 0.4 – 40 | 0.014 | 0.53 | $6.5 \cdot 10^3$ | | [1] |
| AcCHO $\xleftrightarrow{\text{adh}}$ EtOH | 0.3 – 30 | 17 | 0.17 | $1.45 \cdot 10^4$ | | [1] |
| AcCHO $\xleftrightarrow{\text{adh3}}$ EtOH | 0.3 – 30 | 17 | 0.17 | $1.45 \cdot 10^4$ | | [1] |
| $NAD^+ \longleftrightarrow NADH$ | 0.06 – 6 | 0.01 – 10 | 0.01 – 10 | 0.1 | 0.3 – 3 | [2]* |
| $NAD^+ \longleftrightarrow NADH$ | 0.04 – 4 | 0.1 – 100 | 0.001 – 1 | 0.01 | 0.03 – 0.3 | [2]* |

| Michaelis-Menten (Irreversible): $\nu = \frac{V_{\max}}{K_{\mathrm{M}}} \frac{m}{1 + m/K_{\mathrm{M}}}$ | | | |
|---|---|---|---|
| Reaction (enzyme) | $V_{\max}$ ($\mu$mol/min/mg) | $K_{\mathrm{M}}$ $(mM)$ | Citation(s) |
| PYR $\xrightarrow{\text{pdc}}$ AcCHO | 0.03 – 3 | 5 | [1] |
| PYR $\xrightarrow{\text{mcp}}$ PYR | 0.002 – 0.2 | 0.06 – 6 | [3] |
| PYR $\xrightarrow{\text{pdh}}$ AcCoA | 0.01 – 1 | 0.06 – 6 | [4, 5] |

| Inactive Transport (Linear): $\nu = v_{\mathrm{in}} s_{\mathrm{cyto}} - v_{\mathrm{out}} s_{\mathrm{mito}}$ | | | |
|---|---|---|---|
| Reaction (enzyme) | $v_{\mathrm{out}}$ (min$^{-1}$) | $v_{\mathrm{in}}$ (min$^{-1}$) | Citation(s) |
| AcCHO $\leftrightarrow$ AcCHO | $10^{-4} - 1$ | $2 \cdot 10^{-4} - 1$ | |
| EtOH $\leftrightarrow$ EtOH | $10^{-4} - 1$ | $2 \cdot 10^{-4} - 1$ | |

Table S1: Parameters and parameter ranges used in the yeast model. All ranges are taken over a log-normal distribution. Colour indicates where the reaction happens: yellow, blue and yellow to blue for cytosol, mitochondria and cytosol into mitochondria respectively. Additonally, the *pdc* reaction is allosterically regulated by Pyruvate, which is modelled by changing the powers in the Michaelis-Menten reaction (see Fig. S1). Citations to studies used for obtaining parameters are given on the last column of the table. Note that the Pyr $\rightarrow$ Pyr reaction is an active transport reaction, which has the same functional form as an irreversible Michaelis-Menten reaction. In the case of simulations using 'fixed branch' parameters, the parameters for the *mcp* and *pdh* reactions are set to be equal to the parameters of the *pdc* reaction, and allosteric regulation of *pdc* is removed. * For cytosolic and mitochondrial NADH/$NAD^+$, the citation was used for the pool size only.
